# Spatial characterizations of bacterial dynamics for food safety: Modeling for shared water processing environments

**DOI:** 10.1101/2024.11.04.621859

**Authors:** Daniel Munther, Shawn D. Ryan, Chandrasekhar R. Kothapalli, Nerion Zekaj

**Author notes:** Corresponding author: Daniel Munther.

## Abstract

Bacterial dynamics occurring in shared water environments during food processing are typically modeled assuming a homogeneous mixing profile. However, given the tank configurations, and water recirculation and reuse specifications used in many facilities, uniform mixing is not always applicable. Towards this goal, we here developed a novel reaction-diffusion-advection model that captures temporal and spatial variations in the water tanks under dynamic conditions. We utilize the dynamics involved in poultry chilling as an example, as this process features a comprehensive interplay of bacteria, water chemistry and water flow dynamics, as well as determining bacteria levels on carcasses moving into final phases of the food production chain, thus directly influencing public health risk. Well-posedness, existence and uniqueness of positive steady-state solutions with global stability are proved, as well as an estimation of the time scale of convergence to the steady-state solution provided. Simulations are used to verify the analytical results incorporating parameters informed by experimental data from generic, non-pathogenic *E. coli*, and predictively estimate the time to equilibrium. We show that during a typical 8 h processing shift, the model reaches steady state within 2 h, applying this result to validate model simulations against commercial data. The calibrated model predicts a distribution of *E. coli* levels on post-chill carcasses with mean and standard deviation of 3.35 ± 0.56 Log10 CFU/carcass, which closely compares to the experimentally observed distribution of 3.55 ± 0.64 Log10 CFU/carcass in an industrial setting. Our results reinforce the key role of space in quantifying essential mechanisms that govern water chemistry and *E. coli* dynamics during poultry chilling. Our model is an important tool to improve decision making for pathogen control during poultry chilling, as well as a blueprint from which models for processing other commodities like fresh produce and pork can be established.

## 1. Introduction

It has been recently estimated that foodborne disease in the United States may increase to 48 million cases per year [1]. For instance, contaminated leafy greens are estimated to cause over 2 million cases annually [2] and the reporting incidence of illness linked to *Salmonella* in poultry has grown dramatically in the last decade [3]. Government and industry intervention to mitigate this burden has drawn on a variety of tools: intensive data collection at multiple junctures in the supply chain − from regional preharvest field sampling for *E. coli* and *Salmonella* presence on leafy greens [4] to “bio-mapping” *Salmonella* and *Campylobacter* counts through stages of poultry processing plants [5], rapid detection of pathogens [6], utilizing and developing “omics” approaches such as WGA for surveillance [7], as well as GIS systems, big data and risk modeling [8, 9]. Regarding modeling, a recent publication by Allende et al. [10] outlines the pros and cons of several comprehensive QMRA tools for food safety. While these models have significant utility, they still face challenges in terms of accurately describing bacteria transfer at various stages in the supply chain. In fact, this information is crucial for decision making to improve food safety as the WHO reported that 25% of food borne outbreaks are closely tied to cross-contamination events [11].

A crucial driver of cross-contamination concerns shared water environments during food processing [11]. From fresh produce to poultry and pork products, processing steps involving common water exposure span several important commodities. While there is significant literature examining the effect of various interventions (such as temperature and sanitizer efficacy) to controlling pathogen levels during such processing steps, the mechanisms of cross-contamination during these stages are still not well understood [12, 13].

This has prompted focused experimental and modeling studies to elucidate water-mediated pathogen transfer. Regarding poultry processing steps, phenomenological (statistical type) models in [14, 15] and mechanistic type models in the form of ordinary differential equations (ODEs) in are the most current models that have been developed to describe bacterial dynamics. More recent examples have emerged in the fresh produce sector and meat processing chain, such as (statistical) quantification of *Salmonella* cross-contamination during diced tomato flume washing as well as modeling (both ODEs and statistical models) the inactivation of hygiene indicator bacteria during pig carcasses scalding [18]. However, whether explicit (ODEs) or implicit (statistical models), these approaches are based on a uniform mixing assumption regarding water chemistry and bacteria concentration within respective water compartments. Given various water flow dynamics, recent attention to include water reuse and recycling, as well as long tank configurations, the homogeneous mixing perspective may not be appropriate.

In this study, considering water flow and reuse dynamics, we relax the mixing assumption, building a model that can showcase both spatial and temporal variation of bacteria transfer. Primarily, our aim is to develop a model that captures the typical mechanisms involved with processing food items through a shared water step. These include: (i) water mediated cross-contamination, (ii) sanitizer and water chemistry/quality dynamics, and (iii) water flow dynamics, tank configuration, water reuse or recirculation.

As a representative example, we focus on poultry chilling, as this process features a comprehensive interplay of (i) – (iii) above, and because it is a “last step”, determining bacteria levels on carcasses moving into final phases of the food production chain and thus directly influencing public health risk. In particular, the USDA has identified the chilling processing juncture as a critical pathogen control point [19]. Thus, to elucidate and inform decision making that will improve pathogen control during chicken chilling, there is a need for mathematical models capable of describing spatially inhomogeneous dynamics involved with water and sanitizer chemistry and pathogen inactivation. To address this need, we formulate and analyze a novel reaction-diffusion-advection system of the chiller process.

This paper is organized as follows: in Section 2 we discuss model formulation. In Section 3, we prove that the model is well-posed, has a unique positive globally attracting steady-state solution and provides an estimate for characteristic time of convergence to steady state in terms of model parameters. In Section 4, we apply the model to generic *E. coli* contamination dynamics during chicken chilling, validating the model against industrial data as well as illustrating the model properties proved in Section 3. In Section 5, we discuss the importance of spatially explicit models for immersion chilling with counterflow water specifications and highlight future work aimed at understanding influential model parameters for model application towards improved control of human pathogens such as *Salmonella* and *Campylobacter* at the chilling juncture of processing. Finally, we discuss how our model can be adapted to improve food safety decision making for processing other commodities such as fresh produce and pork.

## 2. Model Construction

### 2.1. Dynamics of broiler carcasses and organic load in the chiller tank

**Figure 1** provides an overview of the schematic for chiller tank. Given a tank of length *L* (m), a carcass line speed *s* (carcass·min^-1^), and average mass *m* (kg per carcass), assuming that the birds move along the tank at rate *cp* (m·min^-1^) due to the screw auger motion, and reside in the tank for an average time of *dt* (min), we build the following equation for the density of carcasses *P*(*t, x*) (kg·m^-1^) in the tank at position *x* ∈ [0, *L*] and time *t* ≥ 0:

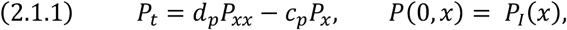

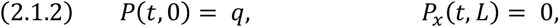

**Figure 1.**
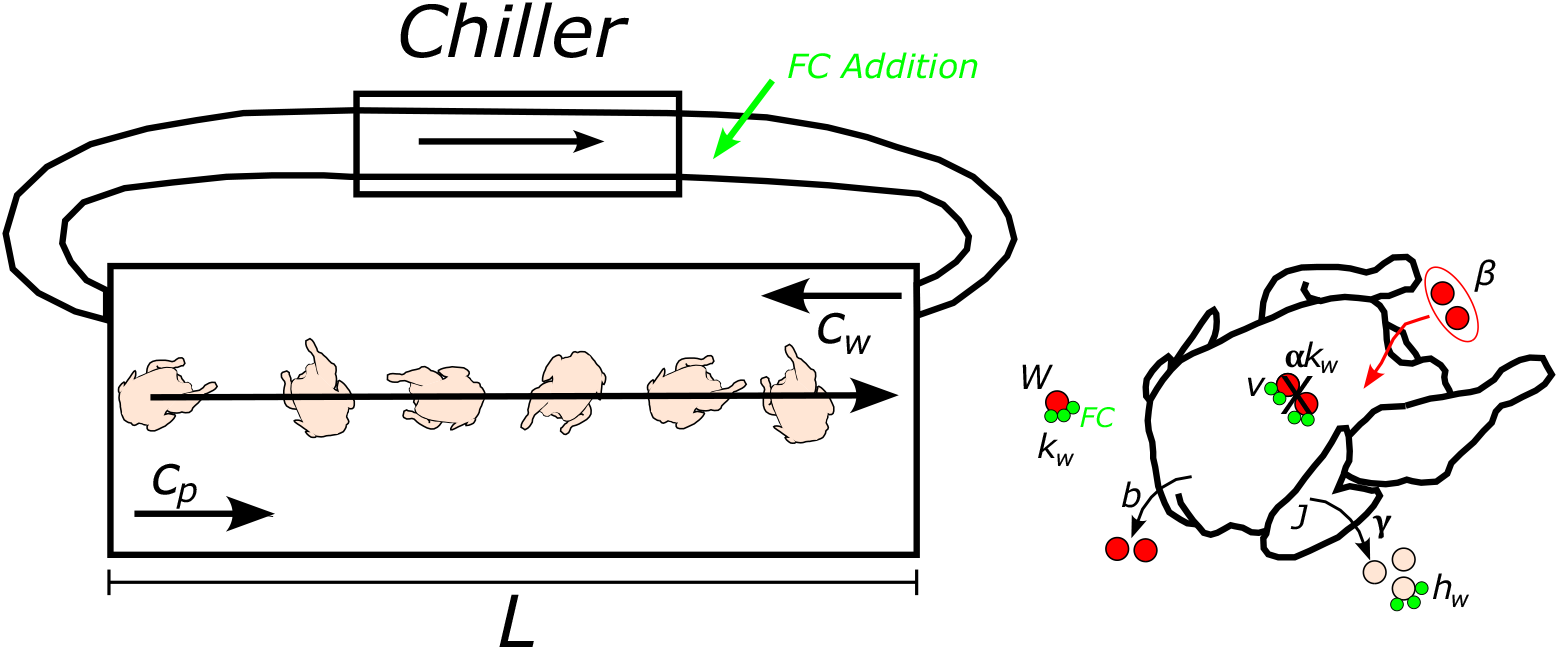
Schematic of the chiller tank. Parameter descriptions can be found in Table 1.

where the position of the birds is also subjected to slight random motion in the tank governed by the diffusion rate *dp* (m^2^·min^-1^) and the boundary conditions indicate a constant density of birds coming into the tank (left boundary condition) and free flow out of the tank (right boundary condition) where 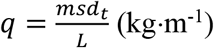. The initial distribution of carcasses in the tank is indicated by *PI*(*x*) (kg·m^-1^).

**Table 1.** Model variables and parameters.

| Name | Description (ref) | Initial value/value | Units |
| --- | --- | --- | --- |
| $P$ | Carcass density in tank (model variable) | $P_I = 0$ | $\text{kg}\cdot\text{m}^{-1}$ |
| $J$ | Organic material on carcass (model variable) | $J_I = 0$ | $\text{kg}\cdot\text{m}^{-1}$ |
| $O$ | Organic material in water (model variable) | $O_I = 0$ | $\text{kg}\cdot\text{m}^{-1}$ |
| $C$ | Free chlorine level in water (model variable) | $C_I = 0$ | $\text{mg}\cdot\text{m}^{-1}$ |
| $v$ | Bacteria level on carcass surface (model variable) | $v_I = 0$ | $\text{CFU}\cdot\text{m}^{-1}$ |
| $W$ | Bacteria level in water (model variable) | $W_I = 0$ | $\text{CFU}\cdot\text{m}^{-1}$ |
| $L$ | Tank length [24] | 9 | m |
| $T_V$ | Tank volume [16] | 50000 | L |
| $d_t$ | Avg. carcass dwell time [24, 33] | 50 | min |
| $c_p = L/d_t$ | Carcass speed thru tank [24, 33] | 0.18 | $\text{m}\cdot\text{min}^{-1}$ |
| $z$ | recirculation rate [24] | 2300 | $\text{L}\cdot\text{min}^{-1}$ |
| $c_w = Lz/T_V$ | Counter flow water speed [24] | 0.41 | $\text{m}\cdot\text{min}^{-1}$ |
| $\gamma$ | Organic load shed rate from carcass, estimated [25] | 0.05 | $\text{min}^{-1}$ |
| $h_w$ | FC depletion rate via material in chiller water [34] | 0.0153 | $\text{m}\cdot\text{kg}^{-1}\cdot\text{min}^{-1}$ |
| $k_w$ | FC kill rate of *E. coli* in water [28] | $0.0127 \pm 0.00029$ | $\text{m}\cdot\text{mg}^{-1}\cdot\text{min}^{-1}$ |
| $\alpha$ | Fraction FC kill rate on carcass surface, estimate [29] | $10^{-4.5 \pm 0.5}$ | No units |
| $b$ | Bacteria shed rate ([24, 32] and refs therein) | $0.05 \pm .0092$ | $\text{min}^{-1}$ |
| $\beta$ | Bacteria attachment rate [31] | $10^{-3.1 \pm 0.5}$ | $\text{m}\cdot\text{kg}^{-1}\cdot\text{min}^{-1}$ |
| $d_p, d_v$ | Diffusion related to carcass [estimated] | 0.004 | $\text{m}^2\cdot\text{min}^{-1}$ |
| $d_o, d_c, d_w$ | Diffusion in tank water [estimated] | 0.004 | $\text{m}^2\cdot\text{min}^{-1}$ |
| $J_o$ | Pre-chill organic material on carcasses [25] | 0.02 | kg/carcass |
| $s$ | Carcass line speed [24] | 144 | Carcass/min |
| $m$ | Carcass mass [estimate -CFIA etc.] | 2 | kg |
| $q = \frac{msd_t}{L}$ | Carcass distribution in tank | 1600 | $\text{kg}\cdot\text{m}^{-1}$ |
| $\sigma$ | Pre-chill *E. coli* load on carcasses [24, 31, 33] | $10^{4.2 \pm 0.5}$ | $\text{CFU}\cdot\text{kg}^{-1}$ |
| $o_b$ | Concentration of organic load in red water recirculated to end of tank (reflects % makeup water added) [estimated based on [34], and addition of fresh water per carcass] | 0.0018 | $\text{kg}\cdot\text{L}^{-1}$ |
| $w_b$ | Fraction of bacterial load in red water recirculated to end of tank | Effectively 0 due to FC dosing at same point in tank. | |

| $c_1$ | Effective FC input concentration [24] | 1.4 | mg·L<sup>-1</sup> |
| --- | --- | --- | --- |

As chickens enter and move through the tank, they release organic material into the chiller water (blood, fat, protein, etc.) which can affect water chemistry as well as microbial counts [20]. We represent the amount of organic material on carcasses at position *x* ∈ [0, *L*] and time *t* ≥ 0 in the tank by *J*(*t, x*) (kg·m^-1^) which satisfies the following:

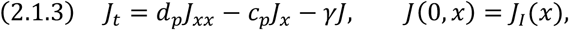

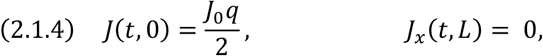

where the dynamics for organic material present at a given moment in time are similar to the carcasses moving through the tank, except we assume that this material enters the water via carcass shedding at an average rate *γ* (min^-1^). Here *J*_0_/2 represents the incoming fraction of organic material to carcass weight (since we assume *m* = 2 kg is the average carcass mass).

Next, let *O*(*t, x*) (kg·m^-1^) be the organic material in the chiller water. To model these dynamics, we assume that fresh water is pumped into the right end (carcass exit) of the tank along with recycled red water taken from the left end (carcass entrance) of the tank. If we let the water recirculation rate be *z* (L·min^-1^) and *T*_*V*_ (L) be the tank volume, then this ensures a countercurrent (relative to carcass direction) water flow speed, 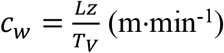 in the chiller tank and leads to the following equations:

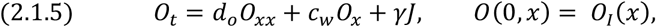

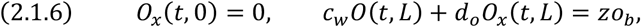

where we assume the organic material in the water diffuses at a rate *d*_*o*_ (m^2^·min^-1^) and is added to the water via carcass shedding at rate *γ* (min^-1^). Notice that the left boundary condition implies the free flow of organic material (with the water pumped out from the tank entrance). This water is chilled and then added to the right side of the tank at rate *z* (L·min^-1^). The right boundary condition, *o*_*b*_ (kg·L^-1^) is the concentration of organic material from the recirculated red water that makes it back into the tank when combined with fresh-water input. While *o*_*b*_ is most likely a function of processing time, for simplicity, we assume that it is constant.

### 2.2. Chlorine kinetics in the chiller water

To track the free chlorine (FC) concentration *C*(*t, x*) (mg·m^-1^) in the chiller water we utilize the following equations:

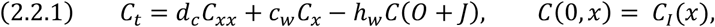

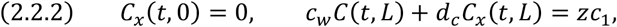

where the diffusion rate is *d*_*c*_ (m^2^·min^-1^), and the flow dynamics (due to water recirculation) are similar to those in equations (2.1.5) and (2.1.6) above. Notice the right boundary condition, *c*_1_ (mg·L^-1^) is the concentration of FC added to the fresh/ recycled water input. The term −*h*_*w*_*C*(*O* + *J*) indicates the rate of FC inactivation via chemical binding with organics in the water and on carcass surfaces. Here, *h*_*w*_ (m·kg^-1^·min^-1^) represents an averaged second order apparent reaction rate of FC with poultry constituents.

### 2.3. Microbial dynamics in the chiller tank

We assume that the following determine the bacteria level *v*(*t, x*) (CFU·m^-1^) on carcasses at position *x* ∈ [0, *L*] and time *t* ≥ 0: (i) bacteria enter the tank on carcasses at a level *σ* (CFU·kg^-1^), (ii) movement of carcasses through the tank, (iii) bacteria shed into the chiller water, (iv) FC inactivation of bacteria on carcass surfaces, and (v) bacteria in chiller water can attach to carcass surfaces. These considerations are captured in the following PDE:

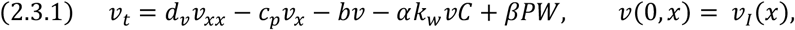

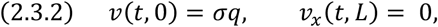

where *b* (min^-1^) is the shed rate, *αk*_*w*_ (m·mg^-1^·min^-1^) is the inactivation rate of bacteria via FC on the carcass surface, *W* (CFU·m^-1^) is the bacteria level in the chiller water, and *β* (m·kg^-1^·min^-1^) is the bacterial attachment rate via water cross-contamination. While the effective contact of FC with carcass surfaces during immersion chilling is complex due to surface morphology affecting various degrees of microbial attachment [21], we take a simplified approach [16]. Specifically, let *k*_*w*_ (m·mg^-1^·min^-1^) be the inactivation rate of bacteria in the chiller water via FC, then we argue that the inactivation rate via FC of microbes on carcass surfaces can be written as *αk*_*w*_, where *α* ∈ (0,1); see Ref [16] for more details.

Finally, we build the following equation governing the dynamics for the bacteria level *W*(*t, x*) (CFU·m^-1^) in the chiller water:

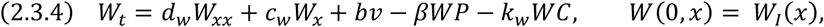

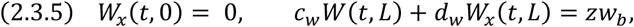

where *W* is subject to the same water recirculation/countercurrent flow as in equations (2.2.1), (2.2.2), (2.3.1), and (2.3.2). Note that *w*_*b*_ (CFU·L^-1^) (in the right boundary condition) is the concentration of bacteria in the recirculated red water which is pumped back into the tank upon being combined with fresh-water input.

### 2.4. Complete model

Combining each of the individual processes described above, we have developed a coupled PDE system for dynamics of the key properties in the poultry chilling process and their associated boundary conditions:

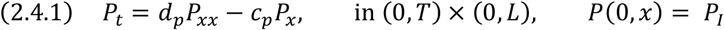

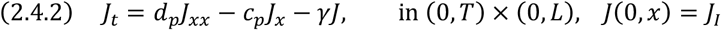

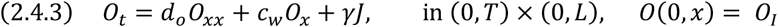

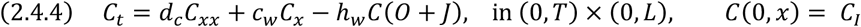

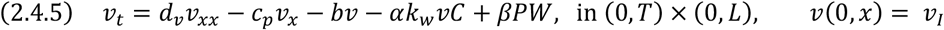

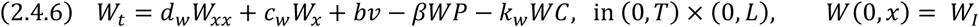

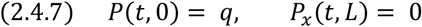

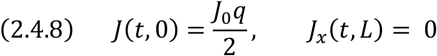

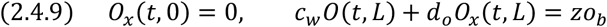

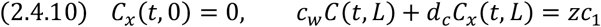

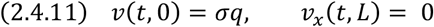

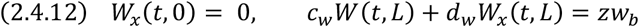

See **Table 1** for a summary of the model variables and parameter definitions.

## 3. Model properties

### 3.1. Well-posedness

To prove well-posedness, we must show that the model equations (2.4.1) − (2.4.6) together with boundary conditions (2.4.7) − (2.4.12) admit a unique positive bounded solution for any set of non-negative initial conditions. To do this, we make use of upper/lower solutions and monotone methods as detailed, for instance, as in Ref [22].

Due to decoupling, we sequentially address equations (2.4.1) – (2.4.4) and (2.4.7) − (2.4.10), respectively, and then treat (2.4.5) − (2.4.6) and (2.4.11) − (2.4.12) as a coupled system. For single equations, we consider the uniformly elliptic operator *L*, defined by

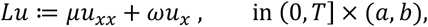

where *µ* > 0, *ω, a, b* ∈ ℝ, with *a* < *b*. Thus, the parabolic boundary value problem with initial condition for a single equation is given as:

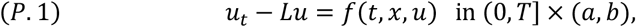

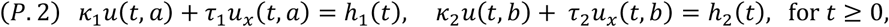

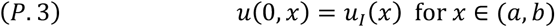

where *κ*_*i*_, *τ*_*i*_ ∈ ℝ and *h*_*i*_(*t*) are smooth for *i* = 1, 2.

For coupled equations, we consider the following parabolic system:

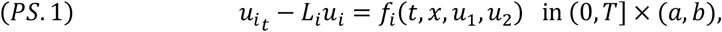

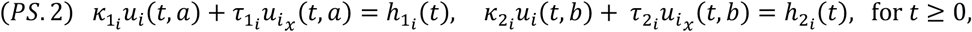

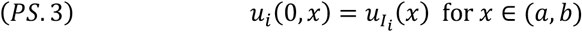

where 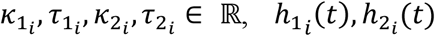 are smooth, and *L*_*i*_*u* ≔ *µ*_*i*_ *u*_*xx*_ + *ω*_*i*_*u*_*x*_ is a uniform elliptic operator for *i* = 1, 2.

Following the theory in [22], if the problem (P.1) − (P.3) has ordered upper and lower solutions û and 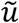 respectively with *f* Lipschitz in *u*, then for any initial condition 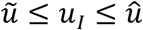, there exists a unique solution that remains between the upper and lower solutions in [0, *T*] × [*a, b*].

#### Theorem 3.1.

*Model (2.4.1)* − *(2.4.6) together with boundary conditions (2.4.7)* − *(2.4.12) has a unique positive bounded solution on* [0, *T*] × [0, *L*] *for any time T* > 0 *and any set of non-negative initial conditions*.

*<u>Proof</u>*. Note that for (2.4.1) and (2.4.2) (with respective boundary conditions), 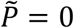 and 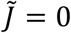 are corresponding lower solutions, while 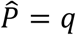 and 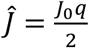 are respective upper solutions. For (2.4.1), *f* = 0, while for equation (2.4.2), *f* = −*γJ*. Thus, for any (*P*_*I*_, *J*_*I*_) in the sector 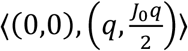, there exists unique, bounded, positive solutions *P* and *J*.

Regarding (2.4.3) and (2.4.9), clearly 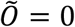 is a lower solution. We claim that 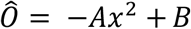 is an upper solution where 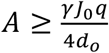 and 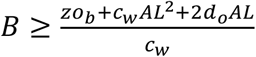. To see this, we have that on (0, *T*) × (0, *L*),

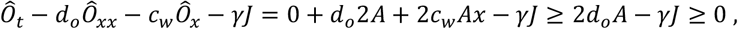

since 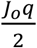 is an upper solution for the equation for *J*. Furthermore, at *x* = *L* we see that for any *t*,

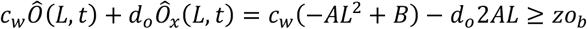

by choice of *B*. Thus, we see that for any 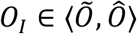, equation (2.4.3) with boundary conditions (2.4.9) has a unique, bounded, positive solution.

For the chlorine equations, (2.4.4) and (2.4.10), again it is clear that 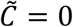 is a lower solution. For the upper solution, we let 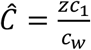. This holds since 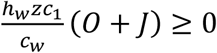 on (0, *T*) × (*a, b*) and since *f* = *h*_*w*_*C*(*O* + *J*) is Lipschitz in *C*.

Finally, we examine (2.4.5) and (2.4.6) with boundary information (2.4.11) and (2.4.12) as a coupled system. (0,0) is a lower solution pair and we claim that 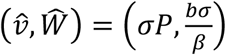 is an upper solution pair, where *P* is any positive solution of (2.4.1) and (2.4.7). For the upper solution pair, consider the following:

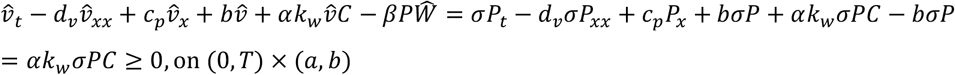

and,

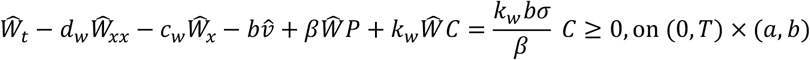

where *C* is any positive solution of (2.4.4) and (2.4.10). Further, note that from equation (2.4.5), *f*_1_(*v, W*) = −*bv* − *αk*_*w*_*vC* + *βPW* and from equation (2.4.6), *f*_2_(*v, W*) = *bv* − *βWP* − *k*_*w*_*WC*. Since 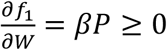 and 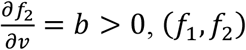, (*f*_1_, *f*_2_) are quasi-monotone non-decreasing.

Furthermore, since *C* is bounded above by 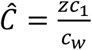 and *P* is bounded above by 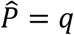, we see that both *f*_1_ and *f*_2_ satisfy the following Lipschitz condition:

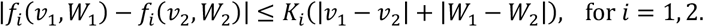

Thus, by monotone theory [22], for any initial conditions in the sector 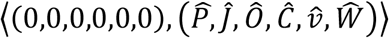, the full model system (2.4.1) − (2.4.6), together with (2.4.7) − (2.4.12), has a unique, positive bounded solution that remains in this sector for all *t* ∈ [0, *T*]. ▪

#### Remark 3.1.1.

*The proof of Theorem 3.1 implies the existence of a uniformly bounded global positive solution, that is, Model (2.4.1)* − *(2.4.6) together with boundary conditions (2.4.7)* − *(2.4.12) has a unique positive uniformly bounded solution on* [0, *∞*) × [0, *L*], *for any set of non-negative initial conditions. This is because the upper solutions bounding each solution component are independent of t and are uniformly bounded on* [0, *L*].

### 3.2. Existence of a unique positive steady state

In this section we consider the steady-state equations for system (2.4.1) − (2.4.6) with (2.4.7) − (2.4.12):

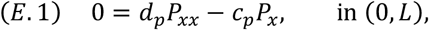

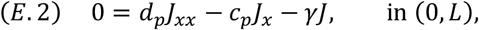

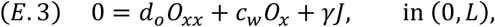

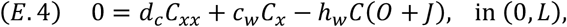

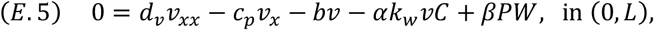

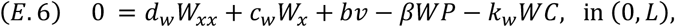

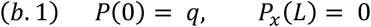

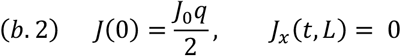

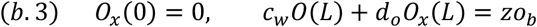

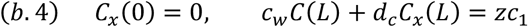

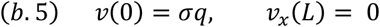

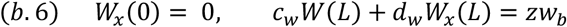

#### Theorem 3.2.

*The steady state system (E.1)* − *(E.6) together with boundary data (b.1)* − *(b.6) has a unique positive solution* (*P*^*^, *J*^*^, *O*^*^, *C*^*^, *v*^*^, *W*^*^).

*<u>Proof</u>*. To demonstrate existence and uniqueness of the positive steady state solution (*P*^*^, *J*^*^, *O*^*^, *C*^*^, *v*^*^, *W*^*^) for system (E.1) − (E.6) with (b.1) − (b.6), we make use of sequential decoupling of the elliptic system.

First, by second order ODE theory, it is clear that *P*^*^ = *q* is the unique positive solution satisfying (E.1) and (b.1). Similarly, for equations (E.2) and (b.2), by second order ODE theory, a unique positive solution *J*^*^ exists.

In addition, by second order ODE theory, equation (E.3) (with *J*^*^ substituted for *J*) with boundary data (b.3) has a unique positive solution *O*^*^ since the corresponding eigenvalue problem

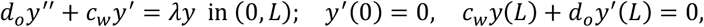

does not have *λ* = 0 as an eigenvalue.

Similarly, since

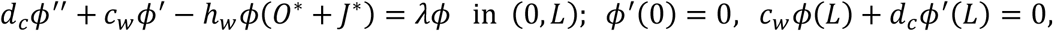

does not have *λ* = 0 as an eigenvalue, by second order ODE theory, equation (E.4) (with *O*^*^ replacing *O* and with *J*^*^ replacing *J*), with boundary information (b.4) has a positive unique solution *C*^*^.

To finish the proof, we focus on the coupled system (E.5) − (E.6) with (b.5) − (b.6) where *P*^*^ = *q* replaces *P*, and *C*^*^ replaces *C*. In this case, it is clear that (0,0) is a lower solution pair and upon inspection one can see that 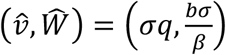 is an upper solution pair for the system. Utilizing monotone theory [22] and the strong maximum principle [23], there exists a solution (*v*^*^, *W*^*^) to (E.5) − (E.6) with (b.5) − (b.6) with *P*^*^ = *q* replacing *P*, and *C*^*^ replacing *C*, such that 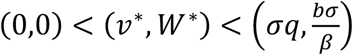. Following monotone theory, there also exist ordered sequences of upper and lower solutions with respective limits 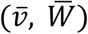 and 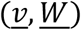 which are themselves solutions of (E.5) − (E.6) with (b.5) − (b.6) and which satisfy:

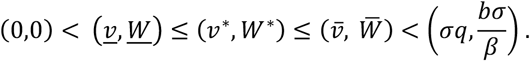

We claim that 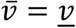 and 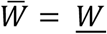. To see this let 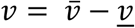 and let 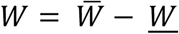. Then by linearity, (*v, W*) satisfy:

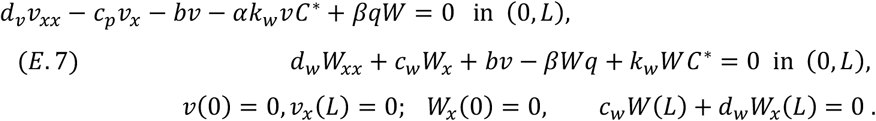

We claim that (*v, W*) = (0,0) is the only solution to (E.7). Clearly (0,0) is a lower solution pair for (E.7). The key idea is to now construct an upper solution sequence that converges to (0,0).

Consider 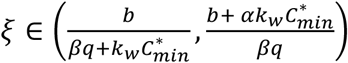, where 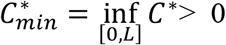. Let *m* be any positive integer and let *k* = *ξm*. Define 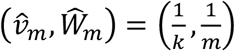, then

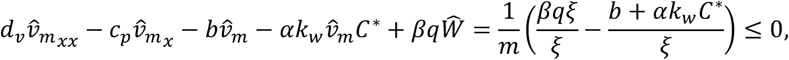

since 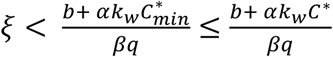.

Also, 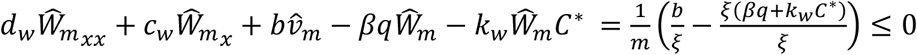, since 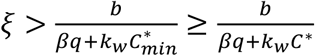. This shows that 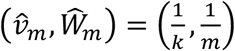, is an upper solution pair and letting *m* → ∞, we have that 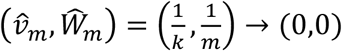. Thus, (0,0) is the unique solution to (E.7), proving that 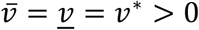 and 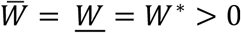 and hence that (*v*^*^, *W*^*^) is the unique positive solution to (E.5) − (E.6) with boundary data (b.5) − (b.6).

Thus, we have shown that (*P*^*^, *J*^*^, *O*^*^, *C*^*^, *v*^*^, *W*^*^) is the unique positive solution of (E.1) − (E.6) with boundary data (b.1) − (b.6). ▪

### 3.3. Global stability of positive steady state

To appropriately apply the monotone theory, we again sequentially decouple the first four equations in system (2.4.1) − (2.4.6), treating single equations separately and then we consider that coupled system (2.4.5) − (2.4.6). Essentially, global stability depends on the uniqueness of the positive steady state (Theorem 3.2) proved in Subsection 3.2 above. For single equations, we make use of the following result from Ref [22]:

#### Lemma 3.3.

(Theorem 4.4 in [22]): *Let* 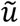, *û be ordered upper and lower solutions of the steady state problem for equations (P.1)* − *(P.2) and let f (in equation (P.1)) be a C*^1^*-function in* 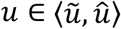. *Then a solution u*^*^ *of the steady state problem for equations (P.1)* − *(P.2) is asymptotically stable with stability region* 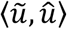 *if and only if it is unique in* 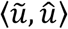.

To treat the remaining, coupled system, we make use of Theorem 5.5 in Ref [22] whose relevant parts are included here for completeness.

#### Lemma 3.4.

(Theorem 5.5 in [22]): *Let* û *and* 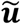 *be an upper solution and lower solution pair, respectively, for the steady state problem of (PS.1)* − *(PS.2). Also suppose that* (*f*_1_, *f*_2_) *is quasimonotone nondecreasing in* 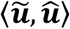. *When the steady state solution* ***u***^*^ *of (PS.1)* − *(PS.2) is unique, then for any initial condition in the sector* 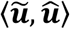, *the solution of (PS.1)* − *(PS.3) converges to* ***u***^*^ *as t* → ∞

These two results lead to the main result of this subsection:

#### Theorem 3.5.

For any non-negative initial conditions, the solution of *model (2.4.1)* − *(2.4.6) together with boundary conditions (2.4.7)* − *(2.4.12) converges to* (*P*^*^, *J*^*^, *O*^*^, *C*^*^, *v*^*^, *W*^*^), *the unique steady state solution of (2.4.1)* − *(2.4.12), as t* → ∞.

*<u>Proof</u>*. For equations (2.4.1), (2.4.7), we apply Lemma 3.3 to lower solution 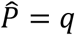 and upper solution 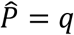 (for the steady state problem), where *f* = 0, and thus have that *P*^*^ = *q* is asymptotically stable in ⟨0, *q*⟩.

For equations (2.4.2) and (2.4.8), we now apply Lemma 3.3 to lower solution 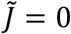 and upper solution 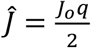 (for the steady state problem), with *f* = −*γJ*, demonstrating that *J*^*^ is asymptotically stable in 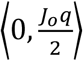.

For equations (2.4.3) and (2.4.9), we invoke Lemma 3.3 with lower solution 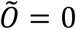 and upper solution *Ô=* Ô*Ax*^2^ + *B* where 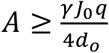 and 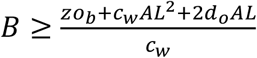, and *f* = *γJ*, to show that *O*^*^ is asymptotically stable in the sector ⟨0, *Ô*⟩.

For the chlorine equations (2.4.4) and (2.4.10), we apply Lemma 3.3 to lower solution 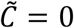 and upper solution 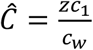, with *f* = *h*_w_*C*(*O* + *J*), demonstrating that *C*^*^ is asymptotically stable in 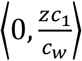.

To complete the proof, we apply Lemma 3.4 to the coupled system (2.4.5) − (2.4.6) with boundary data (2.4.11) − (2.4.12). First, note that from equation (2.4.5), *f*_1_(*v, W*) = −*bv* − *αk vC* + *βPW*, and from equation (M.6), *f*_2_ (*v, W*) = *bv* − *βWP* – *k*_*w*_*WC*. Since 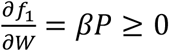 and 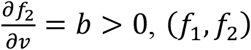, (*f*_1_, *f*_2_) are quasimonotone nondecreasing. Assuming that time has progressed so that *P* = *q* we use the upper solution pair 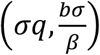 and lower solution pair (0,0) for the steady state problem of (2.4.5) − (2.4.6) with boundary conditions (2.4.11) − (2.4.12). Thus, by Theorem 3.4 we have that for any solution pair (*v, W*), with initial condition in 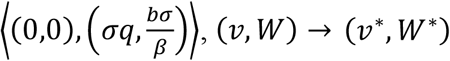 as *t* → ∞.

In conclusion, we have shown that that the basin of attraction of the unique positive steady state (*P*^*^, *J*^*^, *O*^*^, *C*^*^, *v*^*^, *W*^*^) of (2.4.1) − (2.4.6), (2.4.7) − (2.4.12) is 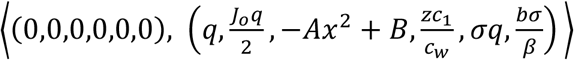, where 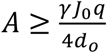 and 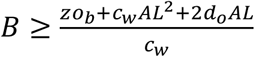.

This can clearly be extended to include any solution with non-negative initial data, and hence we can say that (*P*^*^, *J*^*^, *O*^*^, *C*^*^, *v*^*^, *W*^*^) is globally asymptotically stable for all non-negative initial conditions. ▪

### 3.4. Time scale for convergence to positive steady state

Our goal in this section is to estimate the time for the convergence of system (2.4.1) − (2.4.6), (2.4.7) − (2.4.12) to reach the steady state solution (*P*^*^, *J*^*^, *O*^*^, *C*^*^, *v*^*^, *W*^*^). This will provide a guideline to minimize computational time for running model simulations in terms of predictions as well as parameter estimation.

#### Theorem 3.6.

*For any solution* ***S*** (with nonnegative initial conditions) of (2.4.1)-(2.4.6), (2.4.7)-(2.4.12), ***S***(*T*_*critical*_, *x*) ≈ (*P*^*^, *J*^*^, *O*^*^, *C*^*^, *v*^*^, *W*^*^) on [0, *L*], *where time* 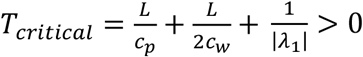 *and λ*_1_ *is the smallest eigenvalue of the matrix* 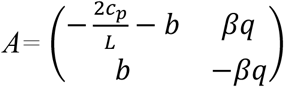.

*<u>Proof</u>*. First, upon inspecting equations (2.4.1) and (2.4.7), we see that *P* converges uniformly to

*P*^*^ = *q* in time 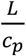. Notice this is the time it takes for the tank to become full of poultry carcasses. In addition, this indicates that a lower bound for convergence time of *J* to *J*^*^ is 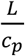. Thus, we assume that for 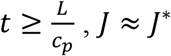, *J* ≈ *J*^*^. Given that *J* is now near steady state, we examine equation (2.4.3) with *J* replaced with *J*^*^:

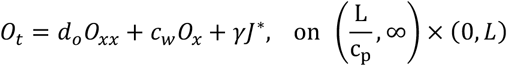

To estimate the time from 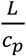 to steady state, we integrate the above equation, and let 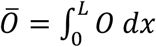, making use of the boundary conditions in equation (2.4.9), to obtain:

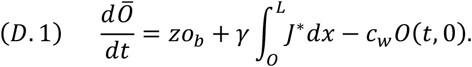

Now, we approximate 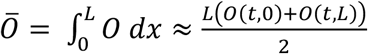, to get that 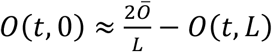. Assuming that *d*_*o*_ ≪ *c*_*w*_ (which is a reasonable assumption as by design the counter flow water speed should be significantly greater than the diffusion rate of organics in the chiller water), from the boundary equation (2.4.9), 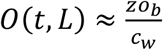. Thus, we have 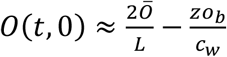. Substituting this into (D.1), we have

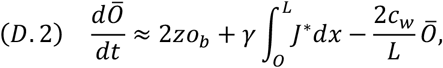

where the solution 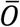 of (D.2) approaches equilibrium in characteristic time 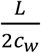.

Thus, our analysis suggests that for 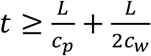, the organic load in the chiller water is near steady state, that is, *O* ≈ *O*^*^. Observe that an increase in the counter-current water speed (*c*_*w*_) will drive the solution toward equilibrium faster.

In terms of the chlorine concentration *C* in the chiller water, we also claim that for 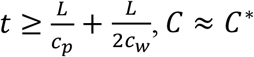. Notice that the inactivation of *C* via organics is given in equation (M.4) by *h*_*w*_*C*(*O*^*^ + *J*^*^). It is reasonable that the characteristic time scale for this reaction, 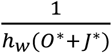 is significantly shorter than 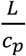, the average dwell time for a carcass to move through tank.

Finally, we separately treat the system for *v* and *W*, equations (2.4.5) − (2.4.6) and (2.4.11) − (2.4.12). Letting 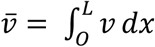 and 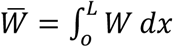, we integrate equations (2.4.5) − (2.4.6) and use the boundary conditions (2.4.11) − (2.4.12) to obtain:

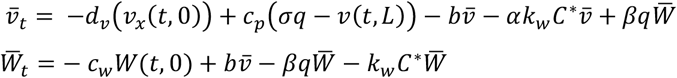

To simplify the above system, we assume that *d*_*v*_ ≈ 0 and 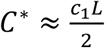. Thus, we have

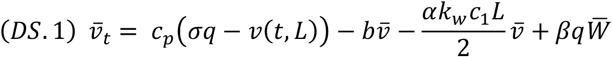

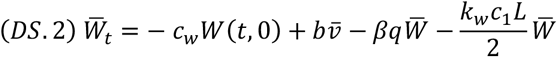

Furthermore, we assume 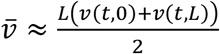 and 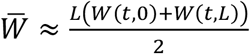. This leads to 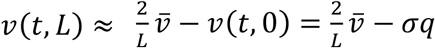 and 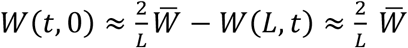, since *W*(*L, t*) ≈ 0 given that the chlorine concentration near *x* = *L* is a maximum value of *c*_1_. Making these substitutions, system (DS.1) − (DS.2) becomes:

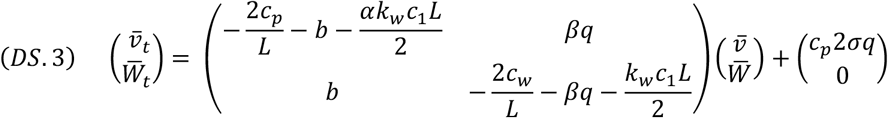

Supposing the bacteria kill rate on the carcass surfaces *αk*_*w*_ ≪ 1, we have that

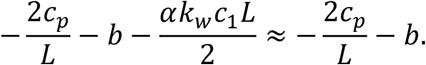

In addition, given that the number of carcasses in the chilling tank *q* ≫ 1, it follows that

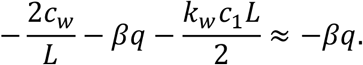

Thus, the matrix for system (DS.3) becomes:

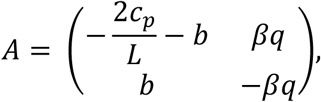

which has eigenvalues 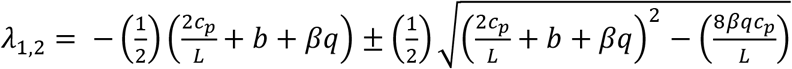.

Notice that Tr(*A*) < 0 and Det(*A*) > 0, indicating that *λ*_1,2_ < 0. Also notice that |*λ*_1_| < |*λ*_2_|. Thus, the slow time scale of convergence of system (DS.3) to steady state is ^1^. ▪

Therefore, we have that the approximate time, *T*_*critical*_, of convergence to the steady state of system (2.4.1) − (2.4.6) with (2.4.7) − (2.4.12) is

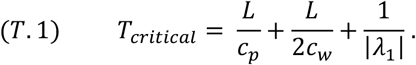

Note that in Section 4, we illustrate that this approximation is quite accurate for the model applied to generic *E. coli* contamination during immersion chilling, using a typical industrial set up as found in Ref [24].

## 4. Model applied to generic *E. coli* contamination during immersion chilling

Using data from the literature as well as our previous modeling studies, we determine parameter values to tailor our model to address the specific dynamics associated to the chiller water chemistry and cross-contamination of broiler carcasses contaminated with non-pathogenic *E. coli*.

### 4.1. Parameter information for organic load dynamics and E. coli contamination

#### 4.1.1 Organic load dynamics

*γ, J*_*o*_, and *o*_*b*_: The organic load in the chiller tank is determined by the line speed *s* (carcass/min), the initial organic load on carcasses entering the tank, *J*_*o*_ (kg), and the shedding of organics from the carcass into the chiller water at rate *γ* (min^-1^). To illustrate the time to convergence estimate, *T*_*critical*_, as well as the steady state solution output (Figure 2), we set *s* = 144 (carcass/min) [24]. Furthermore, we assume a carcass mass on average of *m* = 2 kg and that *J*_*o*_ is 1 % of *m* [25]. Thus, *J*_0_ = 0.02 kg. In addition, from [26], we estimate that *γ* = 0.05 (min^-1^). Finally, we note that 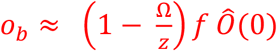, where Ω (L/min) is the addition rate of makeup water per carcass, *z* (L/min) is the water recirculation rate, *f* ∈ (0,1) reflects the fraction of red water return (i.e. post filtering), Ô(0) (kg/L) is the organic load in the chiller water near the tank entrance. Assuming about 3 L of makeup water is continuously added [24], so Ω ≈ 430 (L/min), supposing *f* ≈ 0.65, and Ô(0) ≈ 0.0035 (kg/L) [27], we estimate that o_b_ ≈ 0.0018 (kg/L).

**Figure 2:**
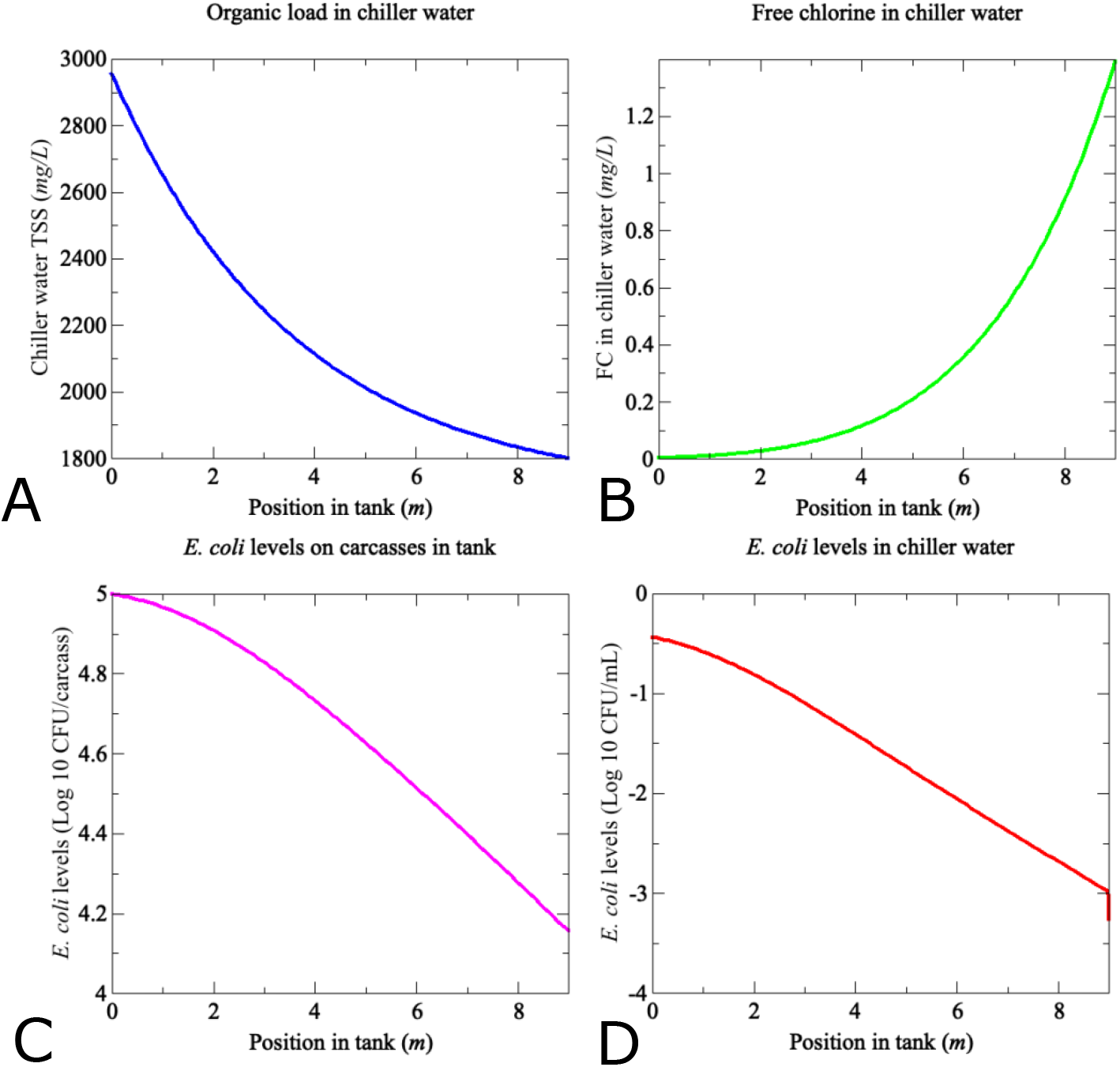
Equilibrium states of model (2.4.1) − (2.4.12) using the parameter values listed in Table 1 (average value if parameter has a range). (Top left) (A) Organic load in chiller water expressed as Total Suspended Solids (TSS) mg·L^-1^, (Top right) (B) Free chlorine level in chiller water (mg·L^-1^), (Bottom left) (C) *E. coli* level on carcasses in the tank (Log10 CFU/carcass), (Bottom right) (D) *E. coli* level in the chiller water (Log 10 CFU·mL^-1^). (ran for *t =* 200 min). Model variables (as presented here) are scaled to units typically utilized in food safety literature. Notice that the weight of carcasses *P* (kg) and organic load material on carcasses *J* (kg) in the tank are not included in the above plots.

#### 4.1.2 FC depletion rate via material in chiller water

*h*_*w*_: From our previous studies [16, 25], we estimated the decay rate of FC due to organic material as 0.0017 (kg^-1^.min^-1^). Scaling this quantity by the tank length, in order to have correct units for the current model, we have *h*_*w*_ = 0.0153 (m.kg^-1^.min^-1^).

#### 4.1.3 FC inactivation kinetics

*k*_*w*_ and *α*: From our previous study [28], the inactivation rate of *E. coli* in water by free chlorine was 70.39 ± 3.19 (L.mg^-1^.min^-1^). Converting this quantity to have the appropriate units for our model, *k*_*w*_ = 0.0127 ± .00029 (m.mg^-1^.min^-1^). While there is a paucity of studies directly measuring *E. coli* inactivation on poultry skin via FC, we use a study by [29] quantifying *Salmonella* inactivation, estimating that 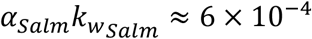. Assuming 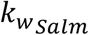 has an approximate order of magnitude of 10^1^ [30], this means that *α*_*Salm*_ ≈ 6 × 10^−5^. From this, we estimate that in the context of *E. coli* inactivation, *α* ∈ 10^[−5,−4]^.

#### 4.1.4. Attachment rate of E. coli to poultry in tank

*β*: Coming from the steady state equation (E.6) with boundary conditions given in equation (b.6), we solve for *β* to obtain 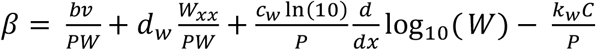. Considering the carcass entrance part of the tank (i.e., *x* ≈ 0), *v* ≈ *σq, P* = *q*, 0 < *d*_*w*_ ≪ 1, *q* ≫ 1, and that 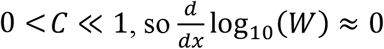, we have then that 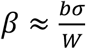. Others [31] found that the *E. coli* level in the red water return ranged from 0.6 to 2 CFU.mL^-1^. From this we assume that *W* ranges in order of magnitude from 0.1 to 1 CFU.mL^-1^ near the carcass entrance of the tank and assuming *σ* = 10^4.75^ CFU/kg (mean of the upper 50% of prechill *E. coli* level on carcass) we estimate that *β* ∈10^[-3.6, -2.6]^.

#### 4.1.5 Shed rate of E. coli from poultry into chiller water

*b*: Assuming an exponential form for bacteria shedding, we have that 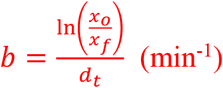, where *x*_*o*_ = initial bacteria concentration, *x*_*f*_ = final bacteria concentration and *d*_*t*_ = 50 min (carcass dwell time in the tank). Following data from [24, 32] and references therein, we suppose a 1-log decrease during chilling, that is 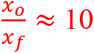. Hence, *b* ≈ 0.05 (min^-1^).

Please refer to **Table 1** for a complete list of model variables, parameter values and associated references.

### 4.2. Application of theoretical results for model validation

Note that the thrust of this section is to portray the model properties proved in Section 3 (Theorems 3.1, 3.2, 3.5 and 3.6); in particular, the features of a typical steady state distribution arising from countercurrent water flow and water recycling dynamics during chilling as well as the reasonableness of utilizing steady state predictions of the model for application during industrial chilling processes.

We showed in Sections 3.2 and 3.3 that model (2.4.1) − (2.4.12) has a unique positive steady state that is globally stable among all non-negative initial conditions (Theorem 3.2 and Theorem 3.5).

While this theory ensures convergence of numerical simulations of the model to a unique solution, Theorem 3.6 also provides us with an estimate on the time for convergence, *T*_*critical*_ from equation (T.1). In this case, 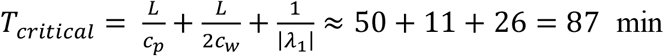, using the parameter values for the water dynamics for the general counterflow chilling process as well as those specific to *E. coli* dynamics (here 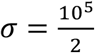, and *α* = 10^−4.5^, *β* = 10^−3^) from Table 1. Simulations were performed in MATLAB R2023b using *pdepe* with space × time mesh *X*_200_ × *TC*, where *X*_200_ = [*x*_0_ *x*_1_ … *x*_198_ *x*_199_], is the mesh along the tank length where *x*_0_ = 0 m, *x*_199_ = 9 m and Δ*x* = *x*_*i*+1_ − *x*_*i*_ = 9/200 m and *TC* = [*t*_0_ *t*_1_ … *t*_139_], is the time step mesh where *t*_*o*_ = 0 min, *t*_139_ = *T* min and Δ*t* = *t*_*i*+1_ − *t*_*i*_ = *T*/140 min. The estimate given by *T* = *T*_*critical*_ is quite good. For instance, using the average parameter values from Table 1, we compared the model output for post-chill *E. coli* levels *v* at time *T* = *T*_*critical*_ ≈ 87 min as well as time *T* = 200 min, finding that the maximum difference in the solutions is given by 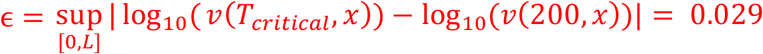 Log_10_CFU/carcass, showing excellent agreement and confirming the time to convergence estimation. **Figure 2** illustrates numerical simulation for *T* = 200 min. (Note that further refinements of the mesh *X*_200_ × *TC* led to negligible error in the solutions and these details are thus not included.)

From a processing viewpoint the three-time components summed to be 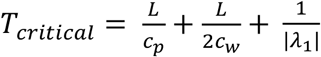also have practical justification. First, 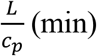 is the time scale for a carcass to travel the entire length of the tank and hence the time needed for the number of carcasses in the tank to equilibrate during a chilling shift. Once this occurs, given the countercurrent water speed *c*_*w*_ (m·min^-1^), the time needed for organic material at the exit end to be transported to the entrance end of the tank is on the order of 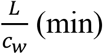. The second term, 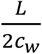 is proportional to this and provides an estimate for the time past 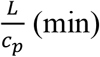 for the organic load in the chiller water to equilibrate. Because the reaction rate with organics is relatively fast, the FC level in the tank essentially equilibrates along with the organic load. The last term, 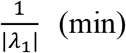 (see the eigenvalue equation in section 3.4) provides an effective time scale balancing carcass speed through the tank against characteristic bacterial shed/attachment time scales.

While bacteria levels and organic load shedding fluctuate per carcass, and as FC input as well as water recycling rates are subject to variability (not to mention bacterial shedding/ attachment dynamics), the model assumes that the distributions associated to these factors change very little on average with respect to processing time. This notion, together with the fact that a typical processing shift is 8 h [24] and by 200 min there is negligible change in the model output, provides mathematical justification to simplify dynamics in the tank and only consider the equilibrium solution.

### 4.3 Model Validation

Others [31] have conducted a study examining the effect of sequential interventions at three commercial poultry processors in the western United States. We apply our model to the immersion chilling stage of “plant B” as described in [31]. In terms of processing specifications, plant B uses a counterflow chiller, red water recycling, a line speed of 275 carcasses.min^-1^, and 20-50 ppm total chlorine input into the chiller water. Testing n=50 pre- and post-chill carcasses, Stopforth et al. [31] determined that the pre-chill *E. coli* distribution was 4.02 ± 0.55 Log10 CFU/carcass (mean ± standard deviation) and the post-chill *E. coli* distribution was 3.56 ± 0.59 Log10 CFU/carcass, where it is assumed that these distributions are both normal.

Using the plant B line speed of 275 carcasses.min^-1^, and assuming 2 kg carcasses each, we set the carcass distribution in the tank *q* = 3056 kg.m^-1^. In addition, using plant B’s data for *E. coli* levels on pre-chilled carcasses, we assume 2*σ* (CFU/carcass) is normally distributed with mean 4.02 and standard deviation 0.55. All the other model parameters are fixed to their corresponding values listed in Table 1 except for the *E. coli* shed rate *b*, the FC kill rate of *E. coli* in water *k*_*w*_, fraction FC kill rate on carcass surface *α*, the *E. coli* attachment rate from water to carcass surface *β*, for which parameter ranges are listed in Table 1.

To explore the variation of these parameters on their respective ranges and the resulting impact on *E. coli* levels of carcasses exiting the chiller tank as well as to validate our model, we used Monte Carlo (MC) simulations. In particular, we simultaneously performed uncertainty analysis and model validation by using Latin Hypercube Sampling (LHS) (n=50) to sample *σ, b, k*_*w*_, *α*, and *β* from their respective ranges listed in Table 1 [35]. Figure 3 illustrates the model prediction (from pre-chill level data to post-chill level predictions) against the post-chill *E. coli* levels observed by [31]. Specifically, the mean and standard deviation of the model predicted distribution for post-chill *E. coli* levels was 3.35 ±0.56 Log10 CFU/carcass compared to the observed distribution with mean and standard deviation of 3.55 ± 0.64 Log10 CFU/carcass.

**Figure 3.**
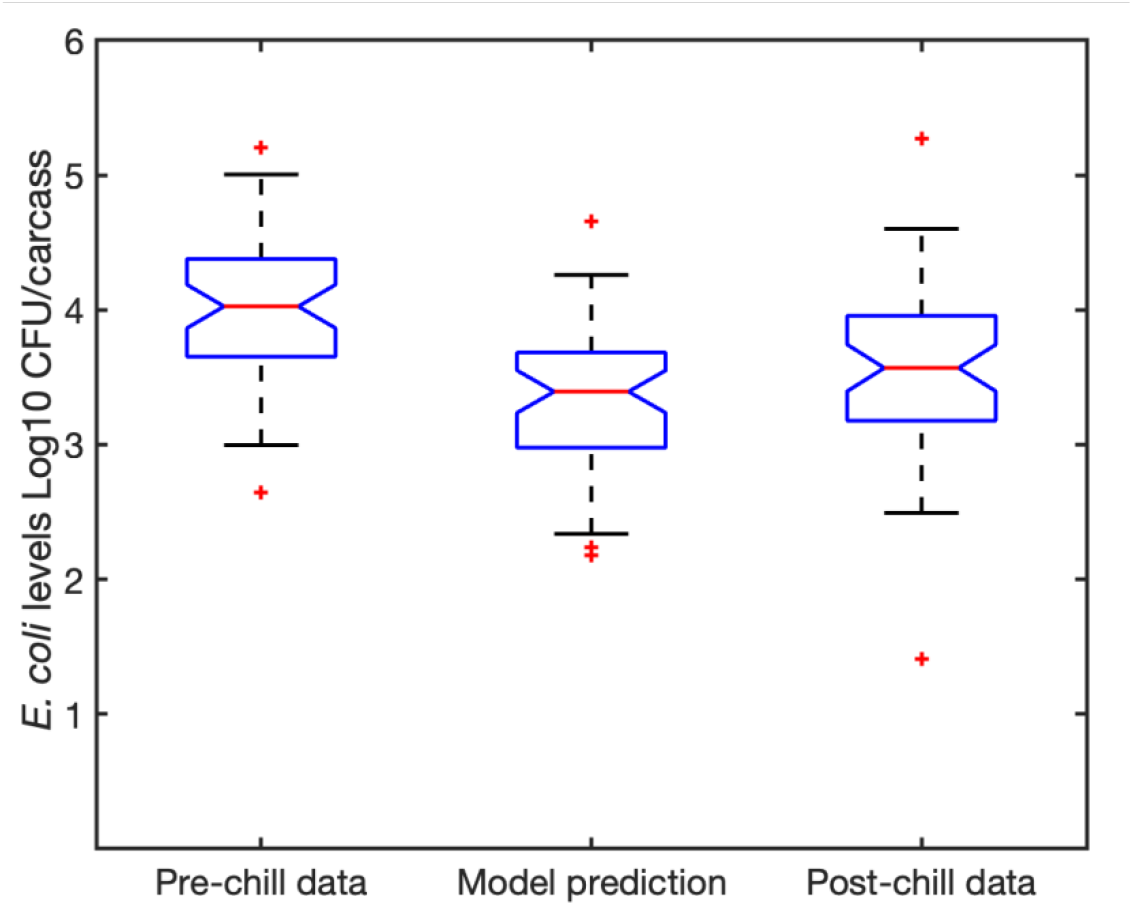
Model performance tested against industrial data from [31]. The pre-chill data corresponds to the *E. coli* level distribution observed in [31] on pre-chill carcasses and, along with parameter values detailed in section 4.3, was used as model input. The model prediction and post-chill data correspond to the *E. coli* level distribution on post-chill carcasses as predicted by the model and as observed in [31], respectively.

Furthermore, we used the two sample Kolmogorov-Smirnov test [36] to evaluate the null hypothesis that the post-chill *E. coli* level data in Figure 3 and the model predicted post-chill *E. coli* levels are from the same continuous distributions. Using the MATLAB function *ks2test*, we find that the test does not reject the null hypothesis at the 0.05 level as it gives a *p*-value of 0.3584 and a test statistic *D*^*^ = 0.18. Notice that a *D*^*^ value close to 0 indicates that the maximum absolute difference between the cumulative distribution functions of the model predicted distribution and the post-chill data distribution is relatively small. This validates that the model describes essential mechanisms dictating bacterial fate during industrial chilling, capturing both average reduction as well as the associated uncertainty on *E. coli* levels of post-chill carcasses observed at the commercial scale.

### 4.4. Effect of countercurrent water flow and water recycling

**Figure 2.** illustrates several key ideas. First, Figure 2 (A) and (B) show how the counterflow and recycling of the chiller water contribute to spatial inhomogeneities in chiller water chemistry. Figure 2 (A) shows a buildup of organic load *O* (mg·L^-1^) in the water at the tank entrance, which decreases along the tank toward the tank exit. When describing the intended effect of a countercurrent chiller at the industrial level, Russell mentions that “a properly operating chiller should have a visible gradient such that the water at the chiller exit is significantly cleaner than the water at the entrance” [20]. This observation matches the model’s steady state prediction for *O* (mg·L^-1^) as the entrance to the tank has a TSS value of about 3000 mg·L^-1^ compared to the exit of the tank where the TSS value is about 1800 mg·L^-1^. Note that the equilibrated chiller water (5 h to 6 h into operation) from a study by Tsai et al. contained about 35% total solids on average [34]. This translates to about 3500 mg·L^-1^ total solids which is slightly higher than the model prediction. Furthermore, when free chlorine is added to the exit end of the chiller, as our model incorporates, Russell indicates that “this approach will result in a ‘clean space’ near the exit end of the chiller. In this space, chlorine will be able to act against bacteria…” [20]. Using parameters from the literature for industrial chilling (Table 1), this phenomenon is illustrated in Figure 2 (B), where in the last 2 m of the tank, the FC level increases from about 0.6 mg·L^-1^ to the effective input level 1.4 mg·L^-1^ (given the input location and organic load distribution) as well as in Figure 2 (C) and (D), where the log reduction of *E. coli* levels on carcasses and in the chiller water is significant in this part of the tank.

Notice that the extent of this portion of the tank, where FC levels are high enough to reduce pathogens, depends on the water recirculation speed, *z* (L/min). Russell indicates that typical heat exchangers are rated for an upper range recirculation rate of about 3600-4000 L.min^-1^ [37] and Morris Re-chiller specifications are such that the recirculation rate can range from about 1500 to 3800 L.min^-1^. To examine the effects of variation in recirculation rates on FC levels and *E. coli* levels on post-chill carcasses, we let z range from 1500 to 4000 L.min^-1^, and again apply our model to the plant B chiller set up as in [31]. To isolate the effect of varying *z*, we adjust the FC input so that the total FC added is the same for each respective case. Specifically, for *z* = 1500 L.min^-1^, we set *c*_1_ = 3.72 mg.L^-1^ and for *z* = 4000 L.min^-1^, we set *c*_1_ = 1.4 mg.L^-1^, where the total effective mass of FC added over 200 min is approximately 1120 g.

**Figure 4** shows the steady state distribution of FC in the tank for both values of *z*. Notice that even though the FC level for *z* = 1500 L.min^-1^ is higher at the exit end of the tank, the slower recirculation speed leads to a larger buildup of organic load in the tank water. Abnavi et al. [28] indicates that even a FC level of 0.25 mg.L^-1^ significantly inactivates *E. coli* in water. From Figure 4, for *z* = 4000 L.min^-1^, *C*(*x*) ≳ 0.25 mg.L^-1^ for *x* ∈ [4,9] m, whereas for *z* = 1500 L.min^-1^, *C*(*x*) ≳ 0.25 mg.L^-1^ for *x* ∈ [6,9] m. Thus, even though the FC level for *z* = 1500 L.min^-1^ is higher at the exit end of the tank (*x* ≈ 9 m), the slower recirculation speed leads to a larger buildup of organic load in the first half of the tank, and hence a smaller portion of the tank in which FC can effectively kill bacteria. **Figure 5** makes this clear as post-chill carcass *E. coli* counts drop on average by about 0.5 Log10 CFU/carcass when only the recirculation rate is increased by about 2.5 times.

**Figure 4.**
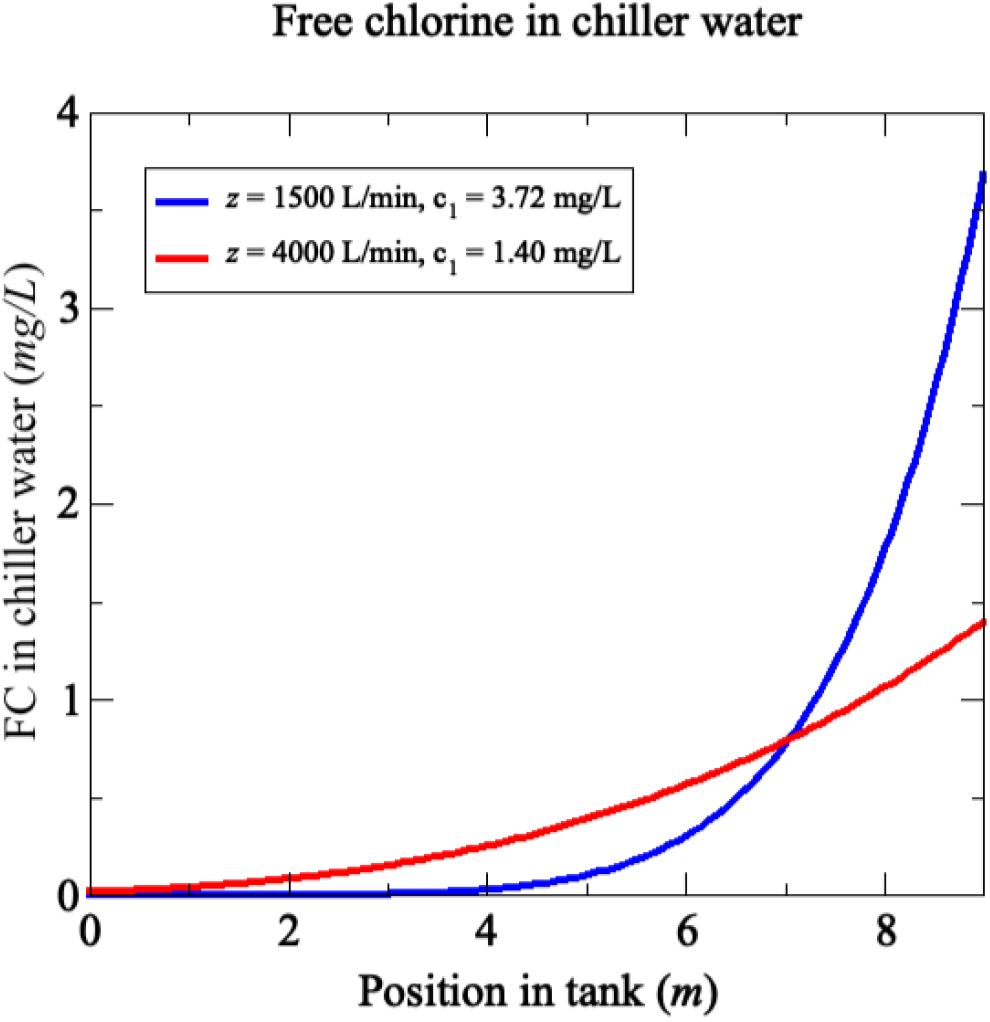
Free chlorine levels in chiller water for low (*z* = 1500 L.min^-1^) and high (*z* = 4000 L.min^-1^) water recirculation rates (other model input parameters were calibrated to the industrial chilling setup in [31] as detailed in section 4.3.). Note that FC input *c*_1_ varies for low and high-water recirculation rates to maintain a total effective mass of FC added over 200 min of approximately 1120 g, respectively.

**Figure 5.**
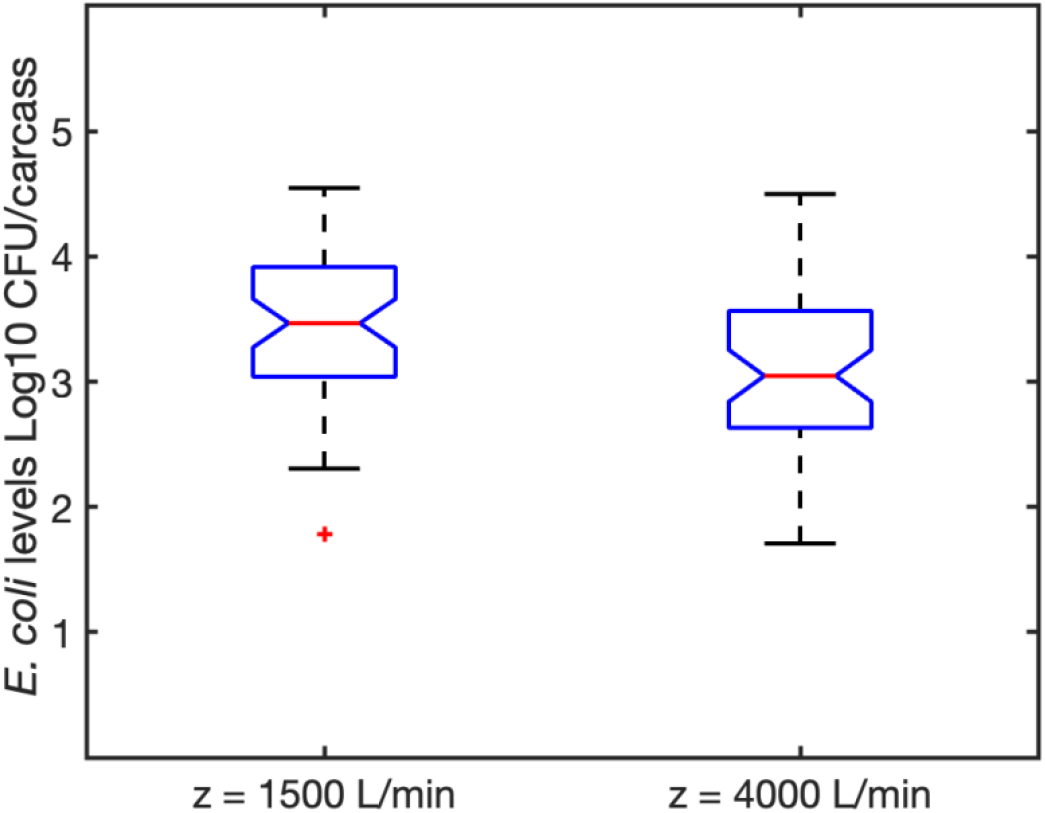
Monte Carlo simulations of *E. coli* levels on post-chill carcasses for low (*z* = 1500 L.min^-1^) and high (*z* = 4000 L.min^-1^) water recirculation rates. Model input parameters were calibrated to the industrial chilling setup in [31] as detailed in section 4.3.

## 5. Discussion

In this work, we developed a novel system of reaction-diffusion-advection equations to quantify bacteria dynamics during a typical industrial chicken chilling process involving water counterflow and recycling dynamics. While these dynamics depend on a complex interplay of water chemistry, cross-contamination, and industrial specifications, our model (2.4.1) − (2.4.12) can quantifiably connect processing control settings and parameters with bacteria levels both in the chiller water and on carcasses surfaces. Our model is armed with the following features: (i) model (2.4.1) − (2.4.12) admits a globally attracting positive steady state solution, and (ii) an expression for the approximate time of convergence to steady state, *T*_*critical*_, expressed in equation (T.1). Our model has an important utility as a tool to improve decision making for pathogen control during poultry chilling as well as a blueprint, in terms of model formulation and analysis, from which models for processing other commodities like fresh produce and pork can be established.

### 5.1. Space is key

While the chilling step is designed to arrest microbial growth on carcasses by lowering their temperature, the water in the tank may still promote significant pathogen cross-contamination if the water−sanitizer chemistry is not controlled during immersion chilling. This is why modern high-speed industrial immersion chiller design incorporates counterflow water dynamics so that the organic content in the chiller water is a decreasing function of tank length [20]. Northcutt et al. observed this phenomenon in an industrial plant using three chiller tanks in series connected to maintain a countercurrent flow of water throughout the process [24]. Examining multiple parameters to represent organic content in the chiller water, from entrance of the first tank to exit of the third tank, they measured a COD change of 1600 mg.L^-1^ to 428 mg·L^-1^, a TSS decrease of 3400 mg·L^-1^ to 975 mg·L^-1^ and a TKN change of 123 mg·L^-1^ to 42 mg·L^-1^, respectively [24]. Coupled with sanitizer delivery at the exit end of the tank, this setup is intended to increase the efficacy of inactivation of bacteria both on carcass surfaces and in chiller water, mostly at the exit end of the tank. Russell emphasizes that this “clean space” feature at the exit end of the tank is the intended effect of the chiller process operating with counterflow water dynamics [20].

From a mathematical standpoint, models addressing mechanisms of bacterial inactivation during chilling range from statistical type models [14, 15] to ordinary differential equation type models [16] to individual based models [25]. A key assumption underlying each of these models is that chemistry changes in the chiller water are independent of space, i.e. tanks are uniformly mixed.

Using a similar mechanistic perspective and mathematical terms as in [16], Sukted et al. developed an ODE model considering 1, 2 or 3 tanks connected in series to ensure water counterflow throughout the chilling process [38]. Their model results showed a similar trend as Northcutt et al. [24] in terms of the decrease in organic load from tank 1 to 3. In comparison with our model (2.4.1) − (2.4.12), the Sukted et al. model is a discrete version, similar to ODE models that treat space as discrete patches. While the Sukted et al. model considers spatial differences among tanks, their model assumes uniform mixing in each tank. Observe that the commercial plant to which they applied their model has tanks that were 6 m, 12 m, and 15 m long, with water flow dynamics of approximately 1900 L·min^-1^ in tank 1, 500 L·min^-1^ in tank 2, and 3200 L·min^-1^ in tank 3. Following the industrial processing specifications under which Northcutt et al. conducted their chiller study, we assumed a countercurrent water flow rate of 2300 L·min^-1^ in model (2.4.1) − (2.4.12) [24] for the steady state simulation in Figure 2.

As Figure 2 illustrates, the well-mixing assumption applied to water chemistry dynamics for tanks 1 and 3 in Sukted et al. model may not be appropriate. Notice, that in our model a key factor that contributes to the spatial inhomogeneity in *O* (mg·L^-1^), and thus the FC level C (mg·L^-1^) (see Figure 2 (A) and (B) at steady state) is the fact that the counterflow water speed, *c*_*w*_ is greater than diffusion rate *d*_*o*_. Therefore, in the presence of water counterflow, to accurately describe the interaction of organic load and sanitizers such as FC, models should explicitly consider space. Figures 4 and 5 further reinforce this notion, demonstrating that bacterial inactivation is a function of counterflow speed even when the total amount of sanitizer added is fixed during a processing period.

Additionally, notice that by examining the equilibrium solution for the system (DS.3) (spatially averaged approximation of the model equations for v and W), we see that from the 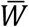 equation, 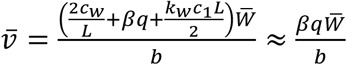 since *q* ≫ 1. Thus, 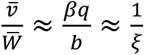, where *ξ* is chosen in Section 3.2. Applying the parameters from Table 1 specific to generic *E. coli* shed and attachment dynamics, we have that 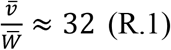 (here both 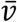 and 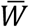 have units CFU). Notice that the numerical calculation of the ratio of the spatial averages of *v*^*^ and *W*^*^ is approximately 90, a similar order of magnitude, providing more justification that the approximate model in (DS.3) captures the bacterial dynamics averaged across the tank quite well. In addition, (R.1) indicates that on average the bacteria levels (total CFU) on the carcasses will not be the same order of magnitude as the bacteria levels in the water (total CFU) and provides some intuition for the choice of *ξ* in Section 3.2 to justify the uniqueness of steady state components for *v* and *W*.

From the spatially averaged perspective, the result in equation (R.1) would seem to be useful from a process control point of view as only bacteria samples from the water (averaging samples taken at the entrance, middle and end of the tank) would need to be obtained to estimate the bacteria levels on carcasses in the chill tank. However, the *spatially explicit situation is more delicate* given water counterflow dynamics. In particular, the ratio of the equilibrium states 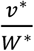, where *v*^*^ (CFU·m^-1^) is the bacteria level on carcasses in the tank and *W*^*^ (CFU·m^-1^) is the bacteria level in the chiller water in the tank, is a function space and is sensitive to the sanitizer concentration. For example, when applied to parameters for *E. coli* and FC sanitization, numerically it can be shown that 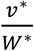 increases from 10^1.6^ to about 10^3.3^, almost 2 orders of magnitude, from the tank entrance (i.e., *x* ≈ 0 m) to the exit end of the tank (i.e., *x* ≈ 9 m). This result corresponds to the FC dynamics as shown in Figure 2 (B) with the increase of FC from almost 0 (mg·L^-1^) at the tank entrance to the input value of 1.4 (mg·L^-1^) at the tank exit. Supporting this notion, if the FC input (*c*_1_ (mg·L^-1^) in Table 1) is set near to 0, the ratio 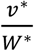 approaches a constant value across the tank. This reemphasizes that mathematical models of chicken chilling processes which utilize sanitizers with counterflow dynamics and intend to accurately capture bacterial inactivation should explicitly consider space.

### 5.2 Future directions

The steady state analysis of model (2.4.1) − (2.4.12), namely the existence of a positive steady state and the time to convergence estimate, *T*_*critical*_, enhance the model’s utility as a food safety tool for poultry chilling. An important future step in this direction concerns the use of detailed spatial data to improve parameter determination as well as to test the sensitivity of model outputs in different locations of the tank. Note that numerically determining specific parameter distributions governing mechanisms of bacterial fate and water chemistry may be computationally expensive given both time and spatial variability. However, as a starting point, model outputs can be fit to spatial data using run times dictated by *T*_*critical*_ to aid with computational efficiency under a wide range of possible parameter values.

In terms of influential parameters, qualitative features of the steady state solutions in Figure 2 suggest a few directions for detailed sensitivity analysis. Figure 2 (C) and (D), imply that during chilling bacteria levels on the chicken as well as those suspended in the chiller water are sensitive to FC levels, which are in turn sensitive to organic load levels in the water (compare Figure 2 (A) and (B)). This implies that the parameter values directly governing the input rates of organic load in the chiller water: for instance, *o*_*b*_ – the fraction of organic load in red water returning to the tank and *γ* - the shed/release rate of organic material from carcasses in the tank, need to be accurately determined. Furthermore, the significant log reduction of *E. coli* on carcass surfaces in the tank (right half of the tank – see Figure 2 (C)) suggests that the FC inactivation rate on the carcass surface (*αk*_*w*_) is important to correctly quantify. Determining the FC inactivation rate as a function of tank location of other bacteria (human pathogens like *Campylobacter* and *Salmonella*) in the context of immersion chilling will be crucial for proper model implementation towards improved decision making for chicken processing safety.

Finally, given the similar dynamics involved with produce washing, we are currently adapting the model development in section 2 and our analysis and applications in sections 3 and 4 to that context. Specific parameter information accounting for the produce/pathogen combination in question will be essential for proper application. In terms of poultry and pork scalding processes, adding a temperature inactivation term (see [26]) to model equations (2.4.5), (2.4.6) involving bacteria changes on carcasses and in process water as well as adjusting boundary conditions to reflect industry practices may provide important insights into optimizing pathogen control.

## Acknowledgments

D.M, S.R., and C.K. acknowledge funding support from the National Institute of Food and Agriculture Grant # OHOW-2022-08999, and the Faculty Research and Development Grant from the Office of Research at Cleveland State University. The authors sincerely thank the two anonymous reviewers for their careful reading and suggestions for improving the manuscript.

## References

[1] A. Chlebicz and K. Ślizewska, Campylobacteriosis, Salmonellosis, Yersiniosis, and Listeriosis as zoonotic foodborne diseases: A review, Int. J. Environ. Res. Public. Health. 15 (2018) 5.

[2] M.R. Powell, Trends in reported illness due to poultry- and nonpoultry associated Salmonella serotypes; United States 1996-2019, Risk. Anal. 44(3) (2024) 641–649.

[3] X. Yang, R. Scharff, Foodborne illnesses from leafy greens in the United States: Attribution, burden, and cost. J. Food. Prot. 87(6) (2024) 100275.

[4] A.M. Belias, A. Sbodio, P. Truchado, D. Weller, J. Pinzon, M. Skots, A. Allende, D. Munther, T. Suslow, M. Wiedmann, R. Ivanek, Effect of weather on the die-off of Escherichia coli and attenuated Salmonella enterica Serovar Typhimurium on preharvest leafy greens following irrigation with contaminated water, Appl. Environ. Microbiol. 86(17) (2020) e00899–20.

[5] D.R. Chavez-Velado, D.A. Vargas, M.X. Sanchez-Plata, Bio-Mapping Salmonella and Campylobacter loads in three commercial broiler processing facilities in the United States to identify strategic intervention points, Foods 13(2) (2024) 180.

[6] F. Fung, H.S. Wang, S. Menon, Food safety in the 21st century, Biomed. J. 41(2) (2018) 88–95.

[7] M.W. Allard, R. Bell, C.M. Ferreira, N. Gonzalez-Escalona, M. Hoffmann, T. Muruvanda, A. Ottesen, P. Ramachandran, E. Reed, S. Sharma, E. Stevens, R. Timme, J. Zheng, E.W. Brown, Genomics of foodborne pathogens for microbial food safety, Curr. Opin. Biotechnol. 49 (2018) 224–229.

[8] J.A. Donaghy, M.D. Danyluk, T. Ross, B. Krishna, J. Farber, Big data impacting dynamic food safety risk management in the food chain, Front. Microbiol. 12 (2021) 668196.

[9] European Food Safety Authority (EFSA), A. F. Jijon, R. Costa, K. Nicova, G. Furnari, Review of the use of GIS in public health and food safety, 19(11) (2022) EN–7639.

[10] A. Allende, S. Bover-Cid, P.S. Fernández, Challenges and opportunities related to the use of innovative modelling approaches and tools for microbiological food safety management, Curr. Opin. Food. Sci. 45 (2022) 100839.

[11] E. Carrasco, A. Morales-Rueda, R.M. Garcia-Gimeno, Cross-contamination and recontamination by Salmonella in foods: A review. Food. Res. Int. 45(2) (2012) 545–556.

[12] K.M. Hardie, M.T. Guerin, A. Ellis, D. Leclair, Associations of processing level variables with Salmonella prevalence and concentration on broiler chicken carcasses and parts in Canada, Prev. Vet. Med. 168 (2019) 39–51.

[13] D. Gombas, Y. Luo, J. Brennan, G. Shergill, R. Petran, R. Walsh, H. Hau, K. Khurana, B. Zomorodi, J. Rosen, R. Varley, K. Deng, Guidelines To Validate Control of Cross-Contamination during Washing of Fresh-Cut Leafy Vegetables, J. Food. Prot. 80(2) (2017) 312–330.

[14] X. Xiao, W. Wang, J. Zhang, M. Liao, H. Yang, W. Fang, Y Li, Modeling the Reduction and Cross-Contamination of Salmonella in Poultry Chilling Process in China, Microorganisms 7(10) (2019) 448.

[15] H. Yang, S. Wang, Y. Li, M.G. Johnson, Predictive models for the survival/death of Campylobacter jejuni and Salmonella Typhimurium in poultry scalding and chilling, J. Food. Sci, 67(5) (2002) 1836–1843.

[16] D. Munther, X. Sun, Y. Xiao, S. Tang, H. Shimozako, J. Wu, B.A. Smith, A. Fazil, Modeling cross-contamination during poultry processing: Dynamics in the chiller tank, Food. Control. 59 (2016) 271–281.

[17] H. Wang, E.T. Ryser, Quantitative transfer and sanitizer inactivation of Salmonella during simulated commercial dicing and conveying of tomatoes, Food. Control. 107 (2020) 106762.

[18] R. A. Stefanello, G. Hermanns, F. Ismael, A. C. Galvao, D. A. Longhi, W. da Silva Robazza, Modeling nonlinear inactivation of hygiene indicator bacteria in pig carcasses during scalding at different pHs, Braz. J. Microbiol (2024) doi: 10.1007/s42770-024-01499-4.

[19] S. J. James, C. James, Chilling and Freezing., in Food Safety Management, Second Edition, Academic Press, 2023, pp. 453–474.

[20] S. M. Russell, Controlling Salmonella in Poultry Production and Processing. CRC Press, 2016.

[21] K.I. Jang, M.G. Kim, S.D. Ha, K.S. Kim, K.H. Lee, D. H. Chung, C.H. Kim, K.Y. Kim, Morphology and adhesion of Campylobacter jejuni to chicken skin under varying conditions, J. Microbiol. Biotechnol. 17(2) (2007) 202–6.

[22] C. V. Pao, Nonlinear Parabolic and Elliptic Equations. Springer, New York, 2013, 777.

[23] M.H. Protter, H. F. Weinberger, Maximum Principles in Differential Equations. Springer, New York, 1984.

[24] J.K. Northcutt, D. Smith, R.I. Huezo, K.D. Ingram,Microbiology of broiler carcasses and chemistry of chiller water as affected by water reuse. Poult. Sci. 87(7) (2008) 1458–63.

[25] Z. McCarthy, B. Smith, A. Fazil, J. Wu, S. D. Ryan, D. Munther, Individual based modeling and analysis of pathogen levels in poultry chilling process, Math. Biosci. 294 (2017) 172–180.

[26] Z. McCarthy, B. Smith, A. Fazil, S.D. Ryan, J. Wu, D. Munther, An individual-carcass model for quantifying bacterial cross-contamination in an industrial three-stage poultry scalding tank, J. Food. Eng. 262 (2019) 142–153.

[27] L.S. Tsai, J.E. Schade, B.T. Molyneux, Chlorination of poultry chiller water: chlorine demand and disinfection efficiency. Poult. Sci. 71(1) (1992) 188–196.

[28] M.D. Abnavi, C. R. Kothapalli, D. Munther, P. Srinivasan, Chlorine inactivation of Escherichia coli O157:H7 in fresh produce wash process: Effectiveness and modeling, Int. J. Food. Microbiol. 356 (2021) 109364.

[29] H. Yang, Y. Li, M.G. Johnson, Survival and death of Salmonella typhimurium and Campylobacter jejuni in processing water and on chicken skin during poultry scalding and chilling, J. Food. Prot. 64(6) (2001) 770–6.

[30] M. D. Abnavi, T. Larimian, P. Srinivasan, D. Munther, C.R. Kothapalli, Inactivation mechanisms of Escherichia coli O157: H7 and Salmonella enterica by free residual chlorine, Environ. Sci.: Water Res. Technol. 8(9) (2022) 2006–2018.

[31] J. D. Stopforth, R. O’Connor, M. Lopes, B. Kottapalli, W. E. Hill, M. Samadpour, Validation of individual and multiple-sequential interventions for reduction of microbial populations during processing of poultry carcasses and parts, J. Food. Prot. 70(6) (2007) 1393–401.

[32] J. K. Northcutt, J. A. Cason, D. P. Smith, R. J. Buhr, D. L. Fletcher, Broiler carcass bacterial counts after immersion chilling using either a low or high volume of water. Poult. Sci. 85(10) (2006) 1802–6.

[33] M.E. Berrang, J.A. Dickens, Presence and level of Campylobacter spp. on broiler carcasses throughout the processing plant. J. Appl. Poult. Res. 9(1) (2000) 43–47.

[34] L.S. Tsai, J.E. Schade, B.T. Molyneux, Chlorination of poultry chiller water: chlorine demand and disinfection efficiency. Poult. Sci. 71(1) (1992) 188–196.

[35] S. Marino, I. B. Hogue, C. J. Ray, D. E. Kirschner, A methodology for performing global uncertainty and sensitivity analysis in systems biology. J. Theor. Biol. 254(1) (2008) 178–96.

[36] J.W. Pratt, J.D. Gibbons, Kolmogorov-Smirnov two-sample tests, in Concepts of Nonparametric Theory, Springer Series in Statistics, 1981, pp. 318–344.

[37] S. M. Russell, Water Reuse in Poultry Processing, in UGA Cooperative Extension Circular, 901 (2023).

[38] N. Sukted, P. Tuitemwong, K. Tuitemwong, W. Poonlapdecha, L.E. Erickson, Inactivation of Campylobacter during immersion chilling of chicken carcasses, J. Food. Eng. 202 (2017) 25–33.

